# RNABag: A Generalizable Transcriptome Foundation Model for Precision Oncology across Biopsy Modalities

**DOI:** 10.64898/2026.04.19.719450

**Authors:** Pengchao Luo, Dong Luo, Dan Li, Xiangyang Xue, Jianbo Yang, Xuejun Gong, Kun Tang

**Affiliations:** HomiGen Intelligence Technology Co., Ltd.; Research Center For Life Sciences Computing, Zhejiang Lab, Hangzhou, 310022, China; Hangzhou Institute for Advanced Study, University of Chinese Academy of sciences, Hangzhou, 310024, Zhejiang, China; The Cancer Center, Union Hospital, Fujian Medical University, Fuzhou, China; Zhejiang Cancer Hospital, Hangzhou Institute of Medicine (HIM), Chinese Academy of Sciences, Hangzhou, 310022, Zhejiang, China; Department of General Surgery, Xiangya Hospital, Central South University, No. 87 Xiangya Road, Changsha, Hunan, 410008, China; National Clinical Research Center for Geriatric Disorders, Xiangya Hospital, Central South University, No. 87 Xiangya Road, Changsha, Hunan, 410008, China; School of Computer and Communication, Jiangsu Vocational College of Electronics and Information, Jiangsu, 223003, China; School of Basic Medical Science, Institute of Molecular Virology andImmunology, Institute of Tropical Medicine,Wenzhou Medical University, 325000, Wenzhou, China

## Abstract

Transcriptomic data is highly sensitive to cancer state and progression, making transcriptome-based foundation models a great promise for diverse clinical ontological inference. However, analyses of transcriptome are conventionally hindered by technical batch effects and limited generalization across platforms. Here, we introduce RNABag, a foundation model designed to generalize well to external datasets. In particular, the model only focuses on highly variable genes to reduce noise; and extensive data augmentation was utilized to pretrain RNABag to learn robust representations, invariant to batch variations. We demonstrate that RNABag achieves superior performance in pan-cancer tissue-of-origin classification and cancer detection in internal validation sets, as well as in zero-shot generalization to external cohorts and in-house clinical samples. Furthermore, RNABag, after specialized finetuning, exhibits strong capabilities in a wide range of clinical applications. The model effectively stratifies patient survival and predicts relapse risks, highlighting key molecular pathways driving tumor progression. Crucially, we extend RNABag’s utility to liquid biopsies, achieving high diagnostic accuracy in plasma cfRNA and tumor-educated platelets (TEPs), thereby supporting its application in non-invasive cancer monitoring. Interpretability analysis revealed pivotal role of tumor immune escape in the cancer induced plasma cfRNA signals. In summary, our study indicates that cancer states and progression may be monitored in details and precision via comprehensive modeling of transcriptome across biopsy modalities.

## Introduction

Early detection, precise molecular subtyping, prognostic risk stratification, and recurrence monitoring are the cornerstones for improving survival outcomes in patients with malignant tumors^1^. In recent years, the deep integration of artificial intelligence (AI) and multi-omics technologies has revolutionized the research paradigm of precision oncology^2^. From intelligent interpretation of digital pathology images^3^ to the mining of tumor-derived signals in circulating cells and nucleic acids^4^, AI-driven tumor detection technologies are rapidly evolving toward multi-omics integration, multi-task joint learning, and pan-cancer coverage, with leaps forward in detection sensitivity, specificity, and clinical applicability.

In computational pathology, foundation models such as CHIEF^5^ and MUSK^6^ have catalyzed a shift toward high-accuracy, pan-cancer evaluation, demonstrating superior performance across dozens of tumor types in tasks ranging from cancer detection, tissue-of-origin (TOO) classification, and prognostic stratification to therapy response prediction. However, pathological images cannot explicitly reflect the molecular signals and gene pathways that form the fundamental basis of tumorigenesis^7^. Meanwhile, pathological testing relies on invasive tissue biopsy, which precludes dynamic and continuous disease monitoring^4^. In our previous study, we proposed a transcriptome foundation model, GeneBag^8^, which demonstrated high sensitivity in cancer diagnosis and prognosis for solid tissue biopsies by leveraging the modeling of full-range RNA expressions to gain profound insights into the gene interaction networks underlying tumorigenesis^8^. Parallel advances in liquid biopsy have sought to enable non-invasive monitoring through circulating analytes. Circulating tumor DNA (ctDNA) based technologies, such as cf-EpiTracing have achieved high sensitivity for multi-cancer detection, disease subtyping, and recurrence and prognosis prediction^9^. Nevertheless, existing cfDNA detection technologies lack sensitivity for early-stage malignancies where tumor-derived DNA is sparse^10,11^. On the other hand, attentions have been paid toward cell-free RNA (cfRNA) and tumor-educated platelets (TEPs). cfRNA provides a high density of functional information, capturing real-time signatures of the tumor microenvironment and systemic immune responses, demonstrating great potentials in early-stage cancer detection, prognosis and treatment response predictions^12,13^. TEPs, acting as systemic “sentinels,” offer an all-in-one platform for pan-cancer detection and TOO identification by altering their RNA repertoire in response to tumor cues^14,15^.

In particular, transcriptomic data is uniquely positioned between genotype and phenotype, offering real-time insights into metabolic reprogramming and immune escape. However, the clinical translation of transcriptome-based AI is stymied by pervasive batch effects and technical noise. Technical variations stemming from library preparation, sequencing depth, and specimen storage conditions often overwhelm biological signals, causing performance drops when models are deployed on independent datasets. Furthermore, existing transcriptomic models are frequently “narrow,” optimized for specific cancers or solid-tissue samples, and lack the robustness required for cross-modality inference in the high-noise environment of peripheral blood.

Here, we present RNABag, a comprehensive transcriptome foundation model designed to monitor the entire continuum of cancer biology across both solid tissues and liquid biopsies. RNABag addresses three critical clinical bottlenecks. First, it mitigates batch effects and generalization barriers by focusing only on highly variable genes (HVGs) and utilizing a rigorous in-silico data augmentation strategy to learn depth-invariant biological patterns. Consequently, RNABag achieves an internal tissue-origin classification accuracy and demonstrates robust zero-shot generalization across external Gene Expression Omnibus (GEO)^16^ and The cBio Cancer Genomics Portal (cBioPortal)^17,18^ cohorts. Second, RNABag extends beyond classification to prognostic evaluation, providing high-fidelity risk stratification for survival and recurrence across 25 malignancies and identifying molecular hubs driving tumor progression via interpretability analysis. Third, we demonstrate that RNABag can successfully decode the complex signals in liquid biopsies, achieving high diagnostic accuracy in plasma cfRNA and TEPs, effectively identifying cancer-induced systemic immune alterations.(https://github.com/DTD007/RNABag)

## Results

### RNABag Architecture and Data augementation

The clinical utility of transcriptomic data based approaches is frequently compromised by pervasive batch effects—stemming from divergent sequencing platforms, protocols, and specimen storage conditions—which impede robust inference across heterogeneous clinical cohorts. To overcome these challenges, we developed RNABag, a Transformer-based foundation model optimized for generalizable cancer representation(Figure. 1A). Unlike its predecessor, GeneBag, which utilized the full transcriptome, RNABag now focuses on 4,096 highly variable genes (HVGs)^19^ derived from the The Cancer Genome Atlas (TCGA)^20^ and Genotype-Tissue Expression (GTEx)^21^ pan-cancer atlases. This targeted approach significantly mitigates computational overhead while filtering transcriptomic noise from lowly expressed or uninformative genes. The architecture comprises eight Transformer^22^ encoder layers with a model dimension of 128 (1.8M total parameters), utilizing the specialized scalar-expression and gene-ID tokenization framework previously described^8^. As delineated in Figure. 1A, the model was trained and validated using a systematic workflow encompassing data augmentation, pretraining, internal validation, and task-specific fine-tuning for survival and recurrence risk prediction, followed by external validation in independent solid tissue cohorts and task-specific fine-tuning for liquid biopsy-based inference. At every stage of this workflow, interpretability analyses were performed to identify key genes and pathways that significantly drive the model’s stratification performance across distinct tasks (Figure. 1A).

**Figure. 1.**
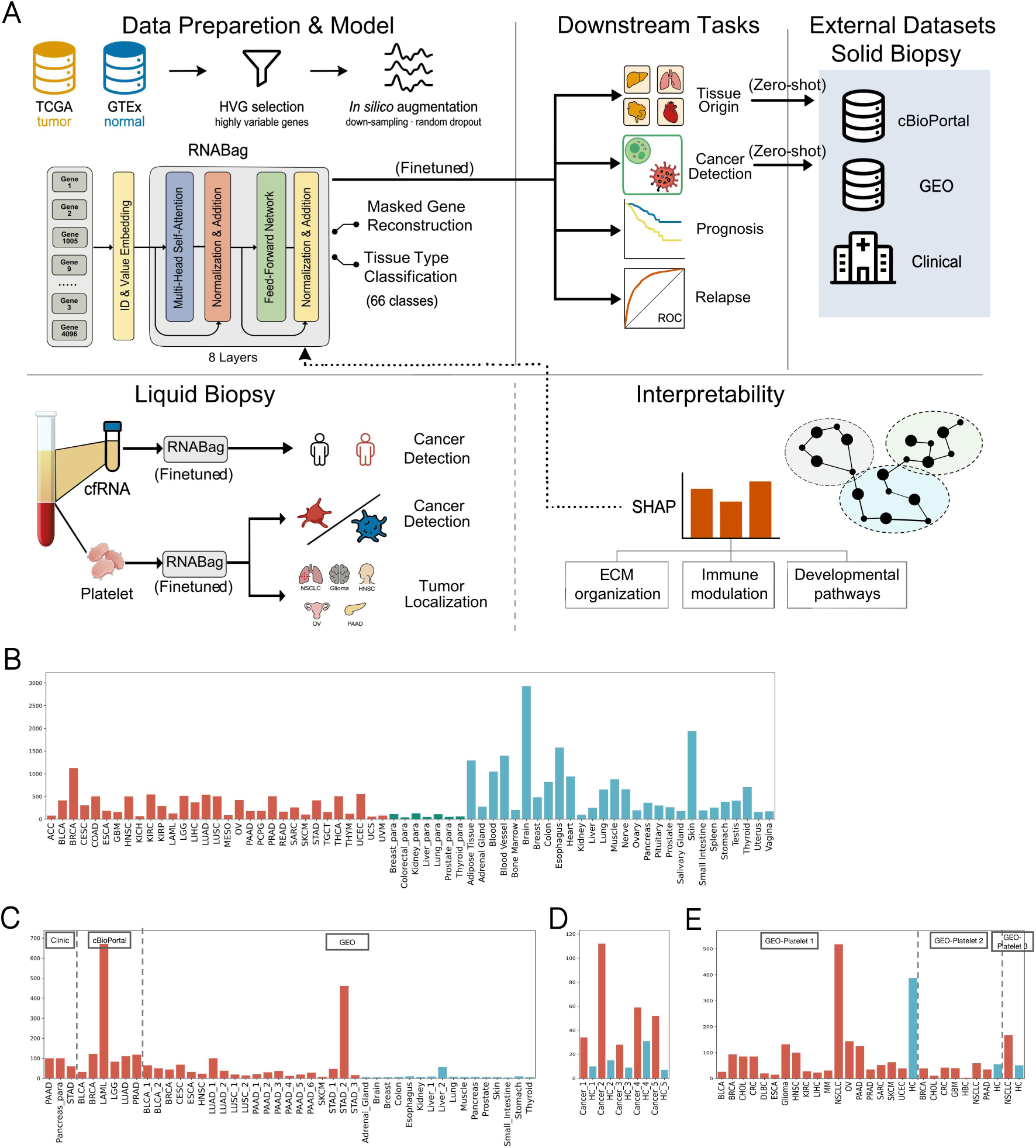
| Schematic of the RNABag framework and dataset characteristics. A, Overview of the architectural design and clinical applications. The RNABag transcriptome foundation model is built upon eight Transformer encoder layers, utilizing multi-head self-attention mechanisms to capture deep gene-gene dependencies. During pretraining, the model integrates large-scale tumor and normal tissue data from TCGA and GTEx. Robust representations are established through highly variable gene (HVG) selection and an in silico data augmentation strategy (such as stochastic down-sampling and dropout), followed by optimization via masked gene reconstruction and multi-class tissue classification tasks. In the downstream transfer phase, the pretrained model demonstrates superior generalization. For solid tissue biopsies, it enables cross-dataset zero-shot identification and cancer detection, and can be fine-tuned for prognostic and recurrence risk assessment. For liquid biopsies, the model effectively adapts to plasma cfRNA and platelet signals, facilitating non-invasive cancer screening and localization. Furthermore, SHAP attribution analysis maps predictive features to key biological pathways, substantially enhancing the biological interpretability of the model’s decisions.B – E, Sample distributions across cohorts. Bar charts detail the sample compositions of the pretraining dataset (B), the independent hold-out validation set (C), the plasma-derived cfRNA cohort (D), and the platelet-derived TEP cohort (E). Red bars indicate tumor samples, whereas cyan bars represent normal or paracancerous tissues. Cancer-type abbreviations follow standard TCGA nomenclature.

TCGA and GTEx datasets form a naturally complementary pair of high-quality pretraining resources, encompassing both tumor and normal tissues. To further mitigate batch effects between these two datasets, we implemented rigorous procedures for data normalization and systematic evaluation of data alignment (Supplementary Figure.1). Furthermore, to account for the substantial difference in sequencing depth between high-quality TCGA/GTEx cohort (30–100M reads/sample) and the external cohorts of routine clinical bulk RNA-seq (5–30M reads/sample), we augmented the training data by generating a series of noise-simulated variants and randomly down-sampling the original profiles to 1/10, 1/20, 1/50, 1/70, 1/100, and 1/280 of their initial depth, coupled with stochastic gene-expression dropout. This training strategy is designed to encourage the model to learn fundamental biological patterns that remain invariant across varying sequencing qualities.

Pretraining followed a dual-task paradigm designed to capture hierarchical biological information. First, a masked gene reconstruction task—where 10% of gene expression values were stochastically masked—required the model to predict continuous expression values based on contextual gene-gene dependencies. Second, a tissue-type classification task challenged the model to accurately identify 66 distinct sample categories across 31 cancer types, 28 normal tissues, and 7 paracancerous tissues(Figure. 1A). RNABag achieved an accuracy of 95.3% in gene reconstruction and 99.6% in tissue classification, substantially outperforming a traditional support vector machine (SVM) baseline under the same evaluation setting (Figure. 2A). To further refine clinical utility, we decoupled malignancy detection from organ identification by training a dedicated tissue-origin classifier, which achieved 99.8% accuracy within the internal validation panel of the TCGA-GTEx cohort, establishing a robust baseline for subsequent zero-shot generalization.

**Figure. 2.**
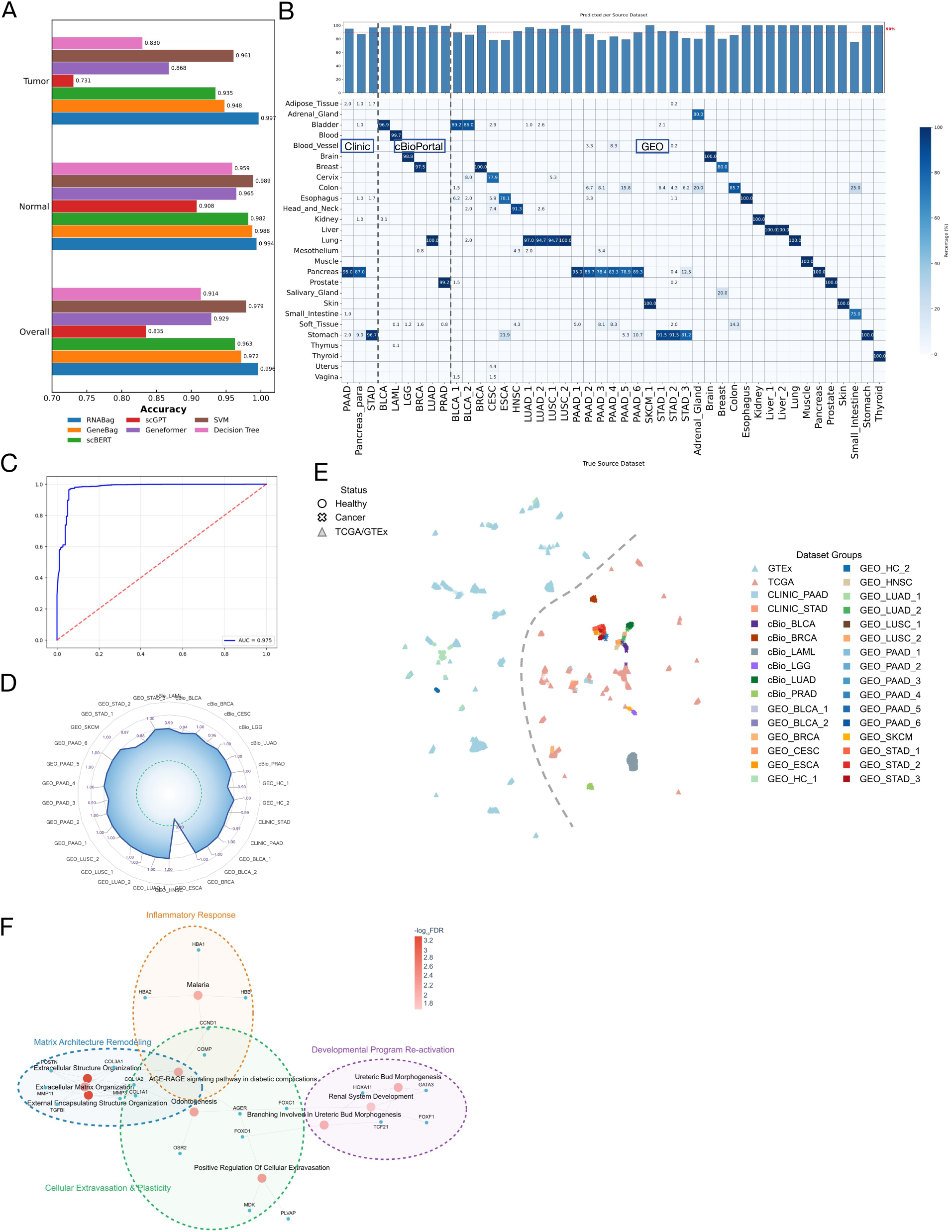
| Multi-task performance evaluation, generalization validation, and biological interpretability of RNABag. A, Comparison of multi-class classification performance. The bar chart demonstrates the accuracy comparison between RNABag and baseline models (including GeneBag, scBERT, scGPT, Geneformer, etc.) across the 66-class task, encompassing Overall, Normal, and Tumor tissue samples. B, Robustness of tissue-of-origin identification. The heatmap and the top bar chart display the model’s tissue-of-origin classification performance on independent hold-out datasets, reflecting its precise identification capabilities within a high-dimensional tissue space. C – D, Validation of cancer detection performance. The ROC curve (C) shows the overall receiver operating characteristic curve for the hold-out datasets in the tumor classification task, achieving an AUC of 0.975. The radar chart (D) illustrates the distribution of recall rates across different subsets of the hold-out datasets. (Note: The C and D labels have been corrected to match the corresponding panels in the provided figure). E, Representation consistency across datasets. The UMAP scatter plot illustrates the spatial distribution of the TCGA-GTEx training set and independent validation sets under the cancer detection model. F, Feature attribution and functional enrichment based on SHAP values. The gene-pathway network diagram displays the top 10 high-confidence pathways and their corresponding key genes in the cancer detection model. Red nodes represent pathways, and blue nodes represent genes. Connecting lines between nodes indicate gene-pathway associations. Color intensity reflects the -log10(FDR) values, showcasing core biological processes such as immune cell regulation and inflammatory responses.

### Zero-shot generalization across external and in-house clinical cohorts

To rigorously evaluate the clinical utility and cross-platform robustness of RNABag, we conducted extensive validation using diverse, external hold-out cohorts (Figure. 1C). We first curated a validation set comprising six bulk RNA-seq datasets from cBioPortal and 22 heterogeneous datasets from the GEO, encompassing 10 cancer types and 15 healthy control (HC) tissues across multiple international clinical centers (Figure. 2B). Despite being entirely unseen during pretraining, RNABag exhibited strong zero-shot generalization in both tissue-of-origin (TOO) classification and malignancy detection. In the TOO task, 16 out of 26 cancer cohorts were correctly assigned to their respective anatomical sites with >90% accuracy (Figure. 2B). Notably, the model demonstrated remarkable consistency across different sequencing centers; for instance, pancreatic adenocarcinoma (PAAD) samples from six independent sources (PAAD_1–6) were accurately identified with recall rates ranging from 78.4% to 95.0%, suggesting that RNABag’s predictions reflect conserved biological signals rather than platform-specific technical noise.

For binary malignancy detection, the model achieved a global Area Under the Curve (AUC) of 0.975 in distinguishing cancerous samples from healthy controls in the external cohorts (Figure. 2C). Stratified analysis confirmed high diagnostic fidelity across heterogeneous cohorts, with 19 of 26 datasets achieving a recall rate of 100% at the default classification threshold (Figure. 2D), Lower performance was observed in specific subsets, such as esophageal carcinoma (ESCA) (0.40) and stomach adenocarcinoma (STAD) (0.87–0.88). The model’s performance was further validated and substantiated using in-house clinical specimens: 60 pathologically confirmed stage I–III stomach adenocarcinoma (STAD) samples from the Affiliated Hospital of Wenzhou Medical University (AHWMU), as well as 100 pathologically confirmed stage I–III pancreatic adenocarcinoma (PAAD) samples paired with 100 adjacent pancreatic paracancerous tissue samples from the Affiliated Hospital of Xiangya School of Medicine (AHXSM). RNABag correctly assigned 95% of PAAD and 96.7% of STAD samples to their anatomical origins, with diagnostic recall rates for malignancy reaching 97% and 95%, respectively (Figure. 2B). A notable observation emerged regarding the transcriptomic identity of paracancerous tissues. Conventionally, these tissues are frequently employed as surrogates for normal controls; yet within the model’s latent space, paracancerous samples formed a distinct cluster clearly separated from both healthy and malignant tissues (Supplementary Figure.2).

Although 87% of AHXSM paracancerous samples were correctly mapped to the pancreas, they demonstrated a distinctive transcriptomic signature that occupied a molecular “middle ground” between physiological homeostasis and overt malignancy, with 34% classified as cancer and the remainder as normal. Such tissues likely harbor early molecular alterations linked to field cancerization, including subclinical inflammatory responses and stromal remodeling. Accordingly, RNABag recognizes paracancerous tissues as a distinct entity and refrains from using them as normal controls.

To confirm that this discriminative power was grounded in meaningful pathophysiology, we employed SHapley Additive exPlanations (SHAP)^23^ to identify the driving molecular features(Figure. 2F). Functional enrichment analysis of top-contributing features revealed a structured, interconnected pathway network defining the pan-cancer landscape, grouped into four core functional modules: extracellular matrix (EM) architecture remodeling, cellular extravasation and plasticity, developmental program reactivation, and inflammatory response. These modules are universally and profoundly dysregulated across solid tumors, underpinning the hallmark malignant phenotypes of invasion, metastasis, and uncontrolled proliferation. Pathways governing extracellular matrix (ECM) synthesis, organization, and remodeling—anchored by core collagen-encoding genes COL1A1, COL1A2, and COL3A1—are known to rewire in malignant cells and cancer-associated fibroblasts, which constitutively activate ECM programs to drive tumor invasion, metastasis, and pro-malignant mechanotransduction signaling^24^; for cellular extravasation and plasticity, the tight constraints on migration and trans-endothelial trafficking in normal mature cells are abolished in malignant cells via aberrant epithelial-mesenchymal transition (EMT) activation^25^, generating highly motile, invasive cells to drive distant metastasis; enrichment of developmental program reactivation reflects a canonical tumorigenesis hallmark, whereby malignant cells hijack latent embryonic proliferation and migration programs to sustain unrestricted growth^26^; finally, the malaria pathway emerges as a significantly enriched signature, driven not by direct malarial infection, but by extensive gene set overlap with inflammatory response pathways, as chronic inflammation is a defining feature of the tumor microenvironment^27^. Importantly, the AGE-RAGE pathway sits at the central nexus of this network, mediating advanced glycation end product (AGE)-dependent cascades regulating ECM remodeling, cellular extravasation, and most prominently the inflammatory response, and the AGE-RAGE axis is a well-validated, high-priority target for anti-cancer drug design and therapeutic development^28^.

### Prognostic Stratification of Pan-Cancer Survival and Recurrence Risks

A central objective of cancer transcriptomics is the high-fidelity prediction of clinical trajectories based on initial diagnostic molecular landscapes. Accurate estimation of overall survival (OS) and the risk of post-therapeutic recurrence is essential for refining personalized treatment, optimizing surveillance protocols, and accelerating the identification of high-value biomarkers. Beyond clinical utility, a fundamental goal is to identify the early, initiating molecular alterations that drive the irreversible invasion-metastasis cascade—the primary biological process underlying post-treatment disease recurrence and incurable metastatic disease. Here, we finetune RNABag to learn shared prognostic signals across all tumor types simultaneously, rather than analyzing individual cancer types separately. This approach not only boosts statistical power through the larger aggregated sample size across diverse malignancies, but may also reveals conserved biological mechanisms underlying malignant metastasis.

In our previous study, we trained the GeneBag model to stratify patients into three discrete survival intervals^8^. Here, we perform more formal survival prediction by implementing Cox partial likelihood loss on a continuous time scale, and further adopt a multi-task joint training strategy to simultaneously predict the death event and gene expression values across datasets for 25 cancer types from the TCGA cohort. The model exhibited robust discriminative performance, with mean the concordance index (C-index) of 0.64 across all in all evaluated malignancies (Figure. 3A). Specifically, RNABag achieved exceptional prognostic accuracy in adrenocortical carcinoma (ACC; C-index = 0.891) and kidney renal papillary cell carcinoma (KIRP; C-index = 0.891), followed by lower-grade glioma (LGG; 0.751), mesothelioma (MESO; 0.739), and acute myeloid leukemia (LAML; 0.69). This prognostic stratification was further supported by Kaplan-Meier analysis, which demonstrated significant divergence between high-risk and low-risk patient groups across multiple cancer types, validated by stringent log-rank tests (Figure. 3B).

**Figure. 3.**
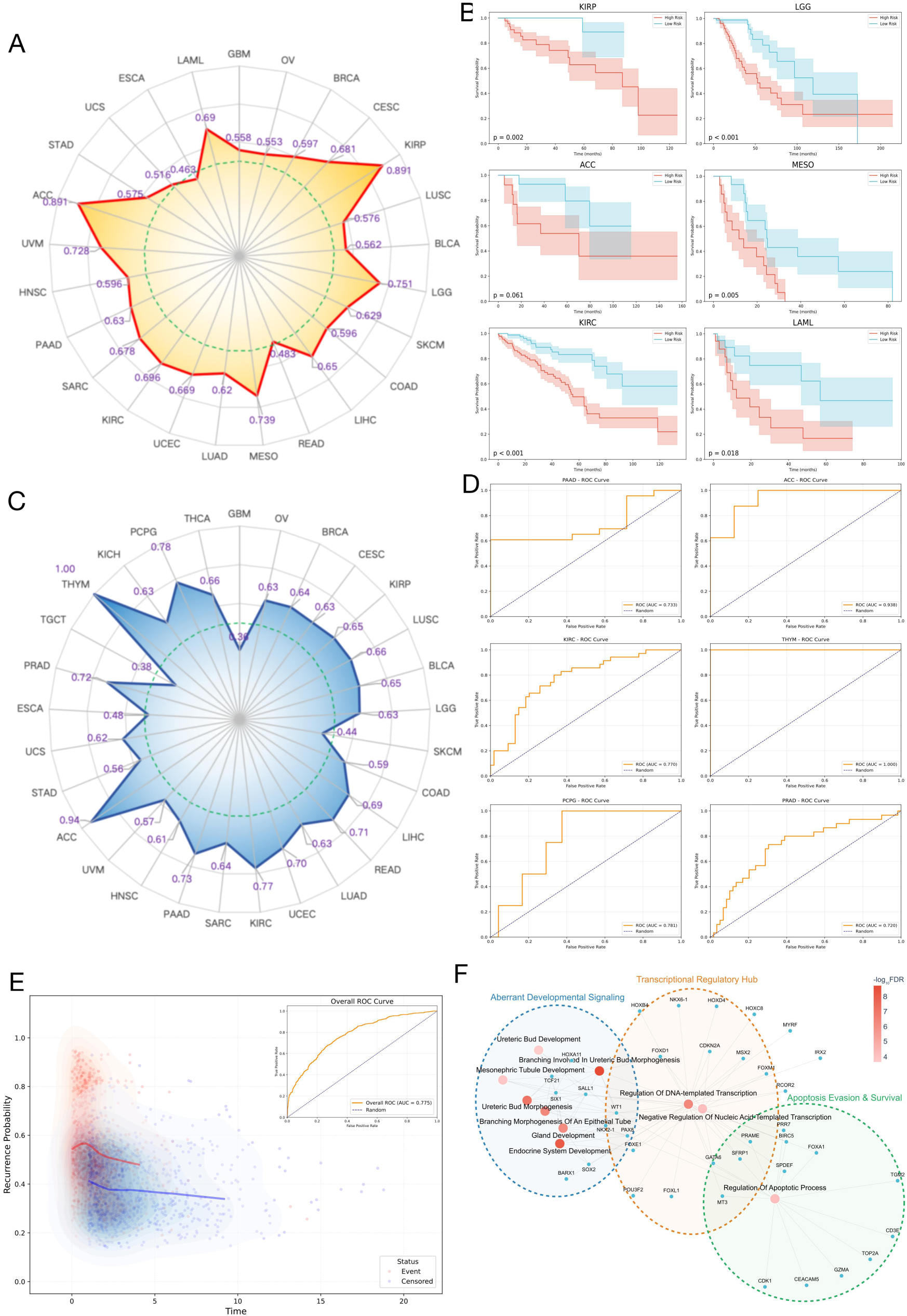
| Performance of RNABag in pan-cancer prognostic survival evaluation and recurrence risk prediction. A, Predictive efficacy of survival analysis. The radar chart illustrates the distribution of the C-index for survival analysis across 25 tumor cohorts, reflecting RNABag’s robust discriminative capacity for patient prognostic risk stratification.B, Survival curves of representative tumors. Kaplan-Meier (K-M) analysis of six cancer types (KIRP, LGG, ACC, MESO, KIRC, and LAML) demonstrates significant survival divergence between the high-risk (red) and low-risk (blue) groups.C – D, Evaluation of recurrence risk prediction. The radar chart (C) systematically presents the AUC for recurrence prediction across 30 tumor types, demonstrating its generalizable performance in capturing long-term clinical outcomes. The receiver operating characteristic (ROC) curves (D) and their corresponding AUC values are displayed for specific cancer types, including PAAD, ACC, KIRC, THYM, PCPG, and PRAD.E, Spatiotemporal distribution of recurrence risk and time. The scatter density plot depicts the temporal trends of recurrence probabilities across the entire cohort. The red curve represents the predicted trend for the event group, whereas the blue curve represents the predictive trajectory for the censored group.F, Elucidation of recurrence-associated molecular mechanisms. The gene-pathway network diagram displays the top 10 high-confidence pathways and their corresponding key genes within the recurrence prediction model. The plot reveals the potential driving roles of biological processes such as transcriptional regulation, endocrine development, and morphogenesis in tumor recurrence.

Beyond survival, we assessed the model’s capacity to predict disease recurrence—a critical hallmark of treatment failure and progression. The recurrence risk prediction task is defined as a binary classification task predicting 12-month post-treatment recurrence risk in pan-cancer patients, to identify high-risk individuals with early short-term recurrence. Training was carried out across 29 cancer types from the TCGA chort. RNABag demonstrated a high overall predictive power (AUC = 0.775) across all cancer types (Figure. 3E). For individual cancer types, the model achieved near-perfect discrimination in thymoma (THYM; AUC = 1.00) and high accuracy in ACC (AUC = 0.94), pheochromocytoma and paraganglioma (PCPG; AUC = 0.78), and kidney renal clear cell carcinoma (KIRC; AUC = 0.77)(Figure. 3D). To access in general, whether this model can predict recurrence, we plotted individualized recurrence probabilities predicted by RNABag and the time-to-recurrence (or time-to-censorship) across the TCGA pan-cancer validation datasets (Figure. 3C).The resultant scatter plot, augmented with kernel density estimates (KDE)^29^, reveals striking differences in the molecular risk landscapes. Patients who experienced a recurrence event (red density cloud) clustered predominantly in regions of high recurrence probability, with a downwards regression line recapitulating a sustained high-risk profile (probability > 0.50), higher at shorter time-to-recurrence and lower at longer time-to-recurrence. Conversely, censored patients populated the lower-probability region (blue density cloud), characterized by a mean predicted probability below 0.40. This separation suggests that RNABag captures clinically relevant risk heterogeneity, supporting its potential for individualized risk stratification.

To elucidate the biological logic underpinning RNABag’s prognosis prediction capacity, we constructed a functional enrichment network of the key features driving recurrence risk (Figure. 3F). This enrichment network identifies three core functional modules significantly associated with pan-cancer postoperative recurrence risk, which directly target the rate-limiting steps of clinical relapse—from minimal residual disease (MRD) survival and cancer stem cell (CSC) maintenance to dormant cell reactivation ^30^. The most prominent module, Aberrant Developmental Signaling (centered on ureteric bud morphogenesis, epithelial branching pathways, and core genes SIX1, SALL1, and HOX family), drives CSC self-renewal, therapy resistance, and reactivation of disseminated dormant tumor cells. These processes are the critical determinants of whether disseminated cells form clinical relapse^31^. The central Transcriptional Regulatory Hub bridges the other two modules, acting as a master upstream switch for tumor cell phenotypic reprogramming. Dysregulation of this module (core genes HOX/FOX families, CDKN2A) enables residual tumor cells to survive postoperative therapeutic and immune pressures, and reinitiate proliferation to drive relapse^32^. The Apoptosis Evasion & Survival module (core genes BIRC5, CDK1, TOP2A) governs MRD survival, the absolute prerequisite for recurrence, allowing residual epithelial tumor cells to escape clearance after surgery and adjuvant therapy^33^.

### Extension of RNABag to Liquid Biopsies for Non-invasive Cancer Monitoring

The clinical utility of transcriptomic foundation models can be significantly expanded by transitioning from solid tissue analysis to the evaluation of circulating analytes. Liquid biopsies, derived from peripheral blood components, offer a minimally invasive and scalable framework for early-onset screening, real-time therapeutic monitoring, and the detection of minimal residual disease (MRD)^4^, thereby bypassing the spatial heterogeneity and procedural risks associated with traditional tissue biopsies^34^. We evaluated the versatility of RNABag by fine-tuning the pretrained model on transcriptomic profiles from plasma cell-free RNA (cfRNA) and tumor-educated platelets (TEPs)^35,36^.

To assess malignancy detection in plasma, we integrated five independent cfRNA cohorts from the GEO database, encompassing 285 malignant samples and 72 healthy controls^37–41^. We fine-tuned the RNABag model pretrained on tissue samples for plasma cfRNA-based cancer detection, appending a plasma-optimized MLP classification decoder and training with cross-entropy loss. Given the restricted sample size, we employed five-fold cross-validation to ensure robust performance estimates. RNABag demonstrated high diagnostic fidelity across all folds, with AUROC values ranging from 0.83 to 0.91 (Figure. 4A). Consistent accuracy (0.79–0.91) and F1 scores (0.88–0.90) across iterations suggest that the model is capable of extracting subtle malignant signals from the high-noise background of the peripheral circulation(Figure. 4B).

**Figure. 4.**
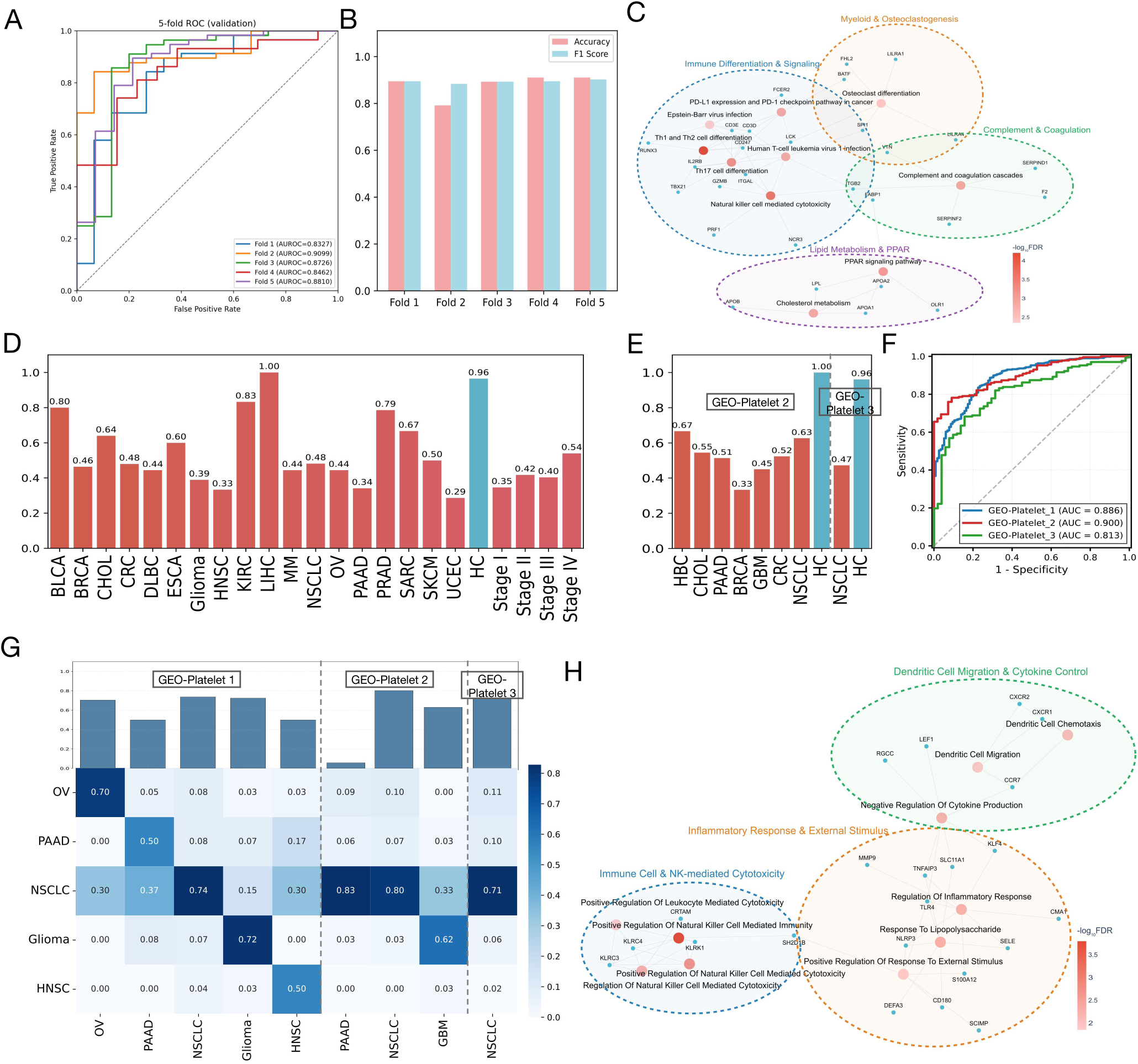
| Application of RNABag in non-invasive cancer diagnosis and tissue tracing based on liquid biopsies. A – C, Diagnostic efficacy and biological representation based on plasma cfRNA. The five-fold cross-validation ROC curves (A) demonstrate the stability of the model in utilizing plasma cell-free RNA (cfRNA) for cancer detection, exhibiting excellent average AUROC performance. The bar chart (B) compares the accuracy and F1 scores across each validation fold. The gene-pathway network diagram (C) reveals the top 10 high-confidence pathways and their corresponding key genes within the cfRNA prediction model, prominently highlighting the immune checkpoint pathway (PD-L1) and various immune cell differentiation processes.D – F, Validation of pan-cancer detection based on tumor-educated platelets (TEPs). The bar chart (D) illustrates the model’s identification accuracy across different anatomical tumor sites and clinical stages (Stage I – IV) within an independent dataset (GEO-Platelet 1). The bar chart (E) displays the classification consistency of the model for diverse malignancies and healthy controls (HC) in the validation sets (GEO-Platelet 2 and 3). The ROC curves (F) systematically evaluate the model’s cancer screening efficacy across the three independent platelet datasets, with AUCs ranging from 0.813 to 0.900.G – H, Tissue-of-origin tracing and functional interpretation of platelet profiles. The confusion matrix heatmap (G) demonstrates the high-resolution performance of the TEP model in predicting the tumor tissue of origin (e.g., OV, PAAD, NSCLC). The gene-pathway network diagram (H) displays the top 10 high-confidence pathways and their corresponding key genes within the platelet prediction model, encompassing biological processes such as natural killer (NK) cell-mediated immune responses, dendritic cell migration, and the regulation of inflammatory signaling.

We further extended RNABag to fine-tune on TEP transcriptome data using an architecture analogous to that employed for plasma cfRNA, analyzing three cohorts encompassing 19 cancer types. Using the largest cohort (GEO-Platelet 1, n=1122) for fine-tuning and two additional cohorts (GEO-Platelet 2 and 3) as independent validation holdouts, the model exhibited high sensitivity in identifying diverse neoplasms^14,15,42^. In the training cohort, RNABag achieved an AUROC of 0.886(Figure. 4D). Notably, at a stringent specificity threshold of 0.965 for healthy controls, the model maintained high recall rates across multiple malignancies, particularly in hepatocellular carcinoma (LIHC, 1.00), kidney renal clear cell carcinoma (KIRC, 0.833), bladder urothelial carcinoma (BLCA, 0.800), and prostate adenocarcinoma (PRAD, 0.786). Analysis across clinical stages revealed an overall upward trend in the model’s performance, identifying 35%, 42%, 40%, and 54% of malignant samples from stages I through IV, respectively at the specificity level of 0.965. Independent validation confirmed this robust efficacy, with AUROCs of 0.908 and 0.810 in GEO-Platelet 2 and 3(Figure. 4F), respectively. Diagnostic performance for CHOL, PAAD, BRCA, GBM, and CRC was consistently replicated. Notably, although hepatobiliary cancer (HBC) was absent from the training set, RNABag achieved a recall rate of 0.67 in the validation holdout at a 100% specificity, suggesting that the model may capture generalized pan-cancer transcriptomic hallmarks beyond cancer-specific signals(Figure. 4E).

Beyond detection, we optimized a dedicated module for TOO prediction in TEPs. RNABag achieved a mean recall rate of 0.63 (range 0.5–0.74), significantly outperforming previously reported benchmarks^15^. This capability for primary site localization was largely preserved in independent holdouts, reaching recall rates of 0.80 and 0.62 for NSCLC and glioma in GEO-Platelet 2, and 0.71 for NSCLC in GEO-Platelet 3. However, we observed that PAAD samples in GEO-Platelet 2 were frequently misclassified as NSCLC (Figure. 4G). This suggests that while TEPs harbor potent diagnostic signals, the organ-specific signatures within the platelet transcriptome may be partially attenuated or convergent across certain epithelial malignancies, necessitating larger, more diverse datasets to refine TOO resolution.

Functional enrichment analysis of the cfRNA cancer classifier revealed four synergistic modules (Figure. 4C). Among these, Immune Differentiation & Signaling is the most prominent module in the network, centered on T helper (Th1/Th2/Th17) differentiation, natural killer (NK) cell-mediated cytotoxicity, PD-L1/PD-1 immune checkpoint signaling, and antiviral immune programs, with core driver genes including CD3D, CD3E, LCK, GZMB, and PRF1. Plasma cfRNA is predominantly derived from circulating immune cells, which undergo systemic transcriptomic remodeling in response to solid tumors even at early stages^40,43^. These pathways represent core host anti-tumor immune responses: malignant transformation drives activation, differentiation, and functional reprogramming of circulating T and NK cells, as well as upregulation of immune checkpoint pathways to mediate immune evasion^44^. The transcriptomic changes in these immune cells are stably released into plasma as cfRNA, making them robust pan-cancer signals for cancer detection.

Similarly, analysis of TEP-derived features revealed that predictive decisions are highly dependent on three core functional modules, including Immune Cell and NK-mediated Cytotoxicity, Inflammatory Response and External Stimulus, and Dendritic Cell Migration & Cytokine Control. All enriched pathways directly reflect the conserved, tumor-induced transcriptomic and functional reprogramming of circulating platelets, which are highly sensitive to even minimal tumor-derived signals (from early-stage solid tumors) and act as central mediators of systemic anti-tumor immunity, cancer-related inflammation, and immune cell trafficking ^45^. Unlike tumor cell-intrinsic signals that are often undetectable in early-stage disease, TEP transcriptomic changes are stable, universal across cancer types, and directly tied to the systemic host response to malignant transformation, which explains their robust significance in pan-cancer analysis.

## Discussion

In this study, we present RNABag, a generalizable transcriptome foundation model engineered to overcome the core barriers of clinical transcriptomic analysis: pervasive batch effects, limited cross-cohort generalizability, and the narrow, task- or disease-specific design of existing analytical frameworks. RNABag enables unified bulk RNA-seq analysis within a single, adaptable foundation model architecture, which is fully fine-tunable for a comprehensive range of oncology tasks, delivers consistent performance across pan-cancer analyses, and flexibly extends to diverse biopsy modalities. Using only TCGA-GTEx cohorts, RNABag learns robust, sequencing depth- and platform-invariant transcriptomic representations via targeted selection of highly variable genes and rigorous in silico data augmentation. We demonstrate robust generalization of RNABag across external clinical cohorts and in-house samples under stringent zero-shot settings. Model performance, generalization, and accuracy can be further substantially improved by incorporating large-scale, diverse real-world clinical data into the base model, either via pretraining data augmentation or fine-tuning tailored to real-world clinical scenarios.

A defining innovation of RNABag is its ability to bridge invasive tissue profiling and non-invasive blood-based monitoring, filling a key gap in existing transcriptomic foundation models. Unlike state-of-the-art models (Geneformer, scGPT) optimized primarily for single-cell data, RNABag’s integrated data augmentation, pre-training and fine-tuning framework is purpose-built to extract meaningful biological signals from heterogeneous clinical data—from high-quality bulk tissue RNA-seq to low-input plasma cfRNA and TEP transcriptomes. Even with limited cfRNA/TEP training data, RNABag achieves high pan-cancer signal sensitivity and cancer type specificity, validating the rich early-tumor information in circulating RNA (via host immune responses)^46^ and the model’s capacity to decode complex tumor-host crosstalk via attention mechanism and transfer learning. Looking forward, pairing transcriptomic foundation models with circulating RNA profiling will enable robust, versatile clinical assays for non-invasive pan-cancer diagnosis and prognosis, supported by large-scale liquid biopsy transcriptomic datasets for systematic model training.

Our interpretability analysis further validates the biological plausibility of RNABag’s predictions, and highlights its potential for translational drug discovery. Across all clinical tasks, the model’s predictive decisions are anchored in well-validated cancer hallmarks, including extracellular matrix remodeling, aberrant reactivation of developmental pathways, systemic immune dysregulation, and inflammatory response. Critically, our analysis independently re-identified 12clinically relevant oncology drug targets: 4 FDA/EMA-approved standard-of-care targets (PD-1/PD-L1 pathway, TOP2A, CDK4/6 axis, coagulation factor F2) and 8 targets in phase I–III clinical development (AGE-RAGE axis, CCND1, BIRC5, PPAR pathway, TLR4, NLRP3 inflammasome, CXCR1/2, MMP9), demonstrating that RNABag can prioritize biologically and therapeutically relevant molecular pathways from complex transcriptomic data. For recurrence risk prediction, the model’s top associated pathways directly target the rate-limiting steps of clinical relapse, including cancer stem cell self-renewal, minimal residual disease survival, and reactivation of dormant disseminated tumor cells, providing mechanistic insights that may inform the development of recurrence-preventing therapies.

In summary, RNABag provides a unified, scalable computational framework for precision oncology across solid tissue and liquid biopsy modalities. Notably, the current model has limitations: constrained per-task training sample sizes and the absence of prospective clinical validation. As such, RNABag at this stage represents only an initial proof-of-concept for the transformative potential of this class of clinical oncology-focused, multi-task transcriptomic foundation models. Future work will focus on expanding training with large-scale real-world clinical datasets, integrating multi-omics data into the framework, validating performance in prospective clinical trials, and extending its utility to guide personalized therapeutic decision-making for patients with cancer.

## Methods

### Data Collection

#### Pre-training Datasets

The pre-training dataset for the RNABag model comprised 29,599 samples spanning 66 tissue types, all of which were acquired from TCGA and GTEx projects through the Recount3 database. The TCGA cohort encompassed 38 tissue types, including 10,014 primary tumor samples across 31 cancer subtypes and 553 adjacent normal tissue samples from 7 tissue types. The GTEx cohort consisted of 19,032 healthy tissue samples covering 28 tissue types and representing major normal anatomical sites throughout the human body, thereby providing a comprehensive transcriptomic foundation for the model to learn the differences between normal and pathological tissues.

### Independent Test Sets

#### cBioPortal Cohort

Six bulk RNA sequencing datasets were curated from the cBioPortal database as components of the independent test set, corresponding to common cancer types across six distinct anatomical sites: bladder urothelial carcinoma, acute myeloid leukemia, lower-grade glioma, breast invasive carcinoma, lung adenocarcinoma, and prostate adenocarcinoma. All datasets were derived from published studies with complete transcriptomic profiles and sample annotation information and were independent of the TCGA and GTEx data, ensuring reliable validation of model performance.

#### GEO Cohort

Twenty-two bulk RNA sequencing datasets were downloaded from the GEO database to form another component of the independent test set. These datasets covered cancerous and normal tissues from 17 anatomical sites, including the adrenal gland, breast, prostate, thyroid, pancreas, esophagus, stomach, small intestine, colon, liver, lung, kidney, bladder, cervix, brain, muscle, and skin, thereby broadening the tissue and cancer-type coverage of the test set and providing diverse data for validating the generalization capability of the model.

#### Clinical Validation Cohort

To validate the practical utility of the model in real-world clinical settings and minimize discrepancies between public-database data and actual clinical data, 260 pathologically confirmed clinical samples were collected from two tertiary medical institutions as an independent clinical validation cohort. These included 100 pancreatic cancer samples and 100 adjacent pancreatic tissue samples from Xiangya Hospital of Central South University, as well as 60 gastric cancer samples from Wenzhou Medical University.

#### Liquid Biopsy Datasets

To extend the application of the RNABag model to non-invasive cancer detection, we incorporated peripheral blood-derived liquid biopsy transcriptomic datasets, including plasma cfRNA and TEPs RNA-seq datasets. All datasets were obtained from published studies with standardized sample collection and sequencing protocols and were used for non-invasive cancer detection and tumor primary-site localization analyses.

#### Plasma cfRNA Datasets

A total of five independent plasma cfRNA sequencing datasets were collected, comprising 357 samples. Each dataset contained 28–112 cancer samples and 7–31 healthy control samples, covering multiple common cancer types. Given the inherent characteristics of plasma cfRNA samples—limited sample size, high technical noise, and low expression abundance—the five datasets were merged for subsequent model training and validation to ensure an adequate sample size and minimize the impact of small-sample bias on model performance.

#### Tumor-Educated Platelet RNA-seq Datasets

Three independent tumor-educated platelet (TEP) RNA-seq datasets from published GEO studies were integrated, totaling 1,496 samples. The first dataset (1,122 samples) comprised 724 cancer samples covering 18 cancer types and 388 healthy controls; the second dataset (284 samples) covered breast cancer, cholangiocarcinoma, non-small cell lung cancer, and other solid tumors with matched healthy controls; and the third dataset (218 samples) included only patients with non-small cell lung cancer and healthy individuals. The three datasets exhibited inherent differences in study origin and sample composition, which were used to evaluate the cross-dataset generalization capability of the model in TEP RNA-seq-based analyses.

### Data Preprocessing and Augmentation

#### Gene Expression Quantification and Normalization

The raw transcriptomic count data from TCGA and GTEx underwent standardized quantification and normalization procedures to reduce technical bias and ensure data consistency. First, raw gene expression counts were converted to fragments per kilobase of transcript per million mapped reads (FPKM)^47^ to minimize the influence of gene length and sequencing depth on gene expression quantification. All subsequent data processing was performed using FPKM values, which unified the quantification standard across different datasets while preserving the biological interpretability of gene expression levels.

#### Highly Variable Gene Selection

To reduce data dimensionality, retain core biological variation in gene expression, and enhance model training efficiency and generalization capability, HVG selection was performed on the quantified and normalized gene set using the scanpy.pp.highly_variable_genes function. The selection process fully considered the expression distribution characteristics of genes in the merged TCGA and GTEx dataset. Ultimately, the 4,096 genes with the highest variability were selected as input features for the RNABag model, ensuring that the selected genes contained key transcriptomic information relevant to tissue characteristics and pathological changes.

#### Data Augmentation

To simulate technical variations in actual sequencing processes, enhance model robustness against random noise, and prevent overfitting, two complementary data augmentation strategies were applied to the preprocessed TCGA-GTEx merged dataset. The augmented dataset was used for model pre-training.

#### Bayesian Downsampling

Bayesian downsampling was performed on the preprocessed gene expression matrix to simulate sequencing depth variations in actual transcriptomic experiments. Samples were generated at six different sequencing depths (1/10, 1/20, 1/50, 1/70, 1/100, and 1/280), covering the common sequencing depth ranges for bulk RNA sequencing. This strategy reduced the model’s over-reliance on specific sequencing depth conditions and enhanced its adaptability to different sequencing data.

#### Random Expression Perturbation

Random expression perturbation was applied to the gene expression matrix to simulate technical noise during sequencing and library preparation. A proportion of gene expression values in the matrix were randomly selected and either set to zero or assigned random values within a low range (0–10). This perturbation strategy disrupted fixed expression patterns for some genes, forcing the model to learn the intrinsic regulatory relationships among genes rather than relying on individual gene expression values, thereby enhancing its robustness to noise.

### Model Architecture

#### Embedding Module

A customized embedding module was designed for the characteristics of bulk RNA sequencing data to achieve deep fusion of discrete genomic identity information and continuous expression abundance signals, thereby serving as the foundation for the RNABag model to capture transcriptomic features. A special CLS token (index 0) was introduced to aggregate the global characteristics of tissue transcriptomes, whereas biological genes were indexed from 1 to 4,096, corresponding one-to-one with the 4,096 highly variable genes selected during preprocessing. Each gene index was mapped to a 32-dimensional identity embedding via one-hot encoding, representing the intrinsic genomic information of the gene; the corresponding FPKM expression value was projected to a 4-dimensional abundance embedding via a linear layer, representing the gene’s transcriptional expression level. The two embeddings were concatenated into a 36-dimensional feature vector, which was then projected to a 128-dimensional fused embedding through a multi-layer perceptron (MLP)^48^, enabling deep feature interaction between gene identity and expression abundance and providing a unified and effective feature representation for subsequent processing by the Transformer encoder.

#### Transformer Encoder

The backbone network of the RNABag model adopted a Transformer encoder architecture to capture long-range gene regulatory dependencies within transcriptomic profiles. The encoder maintained a unified hidden dimension of 128 throughout the network, ensuring feature stability during inter-layer propagation and avoiding information loss due to sudden dimensional changes. It contained 8 stacked encoder layers, each equipped with 8 self-attention heads. The multi-head self-attention mechanism enabled parallel extraction of multi-dimensional long-range gene regulatory features from transcriptomic profiles, fully capturing the intrinsic regulatory associations among different genes. After processing by each layer of the Transformer encoder, the latent state of the CLS token was used as a universal feature descriptor for the entire transcriptomic profile, outputting a 128-dimensional feature vector. This vector served as the core feature representation and was reused as input for all subsequent downstream tasks, achieving unified representation of transcriptomic features across different tasks.

### Pre-training Strategy

#### Input Perturbation and Masking

To ensure that the model learned the intrinsic regulatory relationships among genes rather than relying on the positional information of gene identifiers, input perturbation and masking were applied to the 4,096-dimensional gene feature vectors before they were fed into the pre-training model. In each training epoch, the positions of all 4,096 genes in the input were randomly shuffled to eliminate the model’s dependence on gene order. Simultaneously, 10% of the gene expression values in the feature vectors were randomly selected for masking, with masked positions assigned random values within a low range (0–10), thereby creating a gene expression imputation task that laid the foundation for the model’s multi-task pre-training.

#### Multi-task Objectives

The RNABag model pre-training employed a multi-task supervised learning objective, jointly optimizing two core tasks: gene expression imputation and tissue type classification, enabling comprehensive learning of transcriptomic features. For the gene expression imputation task, mean squared error (MSE) loss was calculated only for the predicted and original values at masked positions to evaluate the model’s ability to reconstruct missing gene expression values and capture fine-grained transcriptomic features. For the tissue type classification task, cross-entropy loss was used for multi-class classification across 66 tissue types to evaluate the model’s ability to recognize and distinguish tissue-specific transcriptomic features. The loss values from both tasks were weighted and summed to obtain the total pre-training loss. Model parameters were updated through gradient backpropagation based on the total loss, enabling the model to simultaneously learn local expression reconstruction and global tissue representation capabilities of transcriptomes.

#### Training Hyperparameters

Pre-training of the RNABag model was implemented in the PyTorch framework using two NVIDIA GeForce RTX 3090 graphics cards to ensure training efficiency. The key hyperparameters were set as follows: a batch size of 2, a maximum of 200 training epochs, the Adam optimizer with a learning rate of 0.0001, and default weight decay to prevent excessive model complexity. During pre-training, the model was evaluated on a validation set (20%) after each training epoch. The Pearson correlation coefficient was used to assess consistency between original and predicted values at masked positions, and classification accuracy was used to evaluate the tissue type classification task, thereby comprehensively validating pre-training effectiveness.

#### Model Evaluation

Based on the pre-trained RNABag model, task-specific models were constructed through transfer learning, and multidimensional, multiscenario evaluations were conducted on independent test sets and clinical validation cohorts.

#### Tissue Origin Classification

The tissue origin classification task aimed to evaluate the model’s ability to accurately distinguish tissue- and organ-specific transcriptomic regulatory features from technical noise and pathological variation. The test set consisted of a multicenter, cross-platform composite dataset integrating 6 cBioPortal cohorts, 22 GEO datasets, and 260 clinical samples, covering cancerous and normal tissues from major anatomical sites throughout the human body and fully testing the model’s feature-recognition capability and robustness to batch effects. The pre-trained RNABag encoder was used as the feature-extraction backbone, with its parameters jointly fine-tuned with a task-specific MLP decoder on downstream-task data. The MLP decoder mapped the 128-dimensional feature vector output by the encoder to discrete tissue-origin labels. Confusion matrices were used to quantify classification performance by calculating precision, recall, and F1 score for each tissue type. The average F1 score across all tissue types was used as the core evaluation metric for comparison with other mainstream models, with a focus on tissue-origin recognition capability and robustness to batch effects in multisource, cross-platform data.

#### Tissue-Based Cancer Detection

The cancer detection task for tissue samples was designed as a binary classification task to distinguish tumor tissues from matched normal tissues at the same anatomical site, aiming to validate whether the transcriptomic features extracted by the RNABag model contained sufficient pathology-related signals to accurately identify expression changes induced by malignant transformation. The tissue-sample cancer detection model was built on the embedding module and encoder of the pre-trained RNABag model, followed by a single-hidden-layer MLP decoder that output the probability of a sample being malignant through a softmax function. During task-specific training, the embedding module and encoder were unfrozen and fine-tuned jointly with the classification decoder on the TCGA-GTEx merged dataset. This task employed a cross-center independent validation strategy: the TCGA-GTEx merged dataset was used as the training set and internal validation set for fine-tuning (stratified random split at 80%/20%). After fine-tuning, all 6 cohorts from cBioPortal, 22 datasets from GEO, and 160 clinical samples were used as independent external test sets without participating in parameter updates or hyperparameter selection, thereby strictly evaluating the model’s generalization performance in real-world clinical scenarios. Evaluation metrics included the AUROC, precision, and recall, comprehensively reflecting the model’s diagnostic capability across different prevalence distributions. AUROC served as the key comparative metric for validating the advantage of the RNABag model over other mainstream models in tissue-sample cancer detection tasks.

#### Survival Outcome Prediction

The survival outcome prediction task aimed to evaluate the model’s ability to perform survival-risk stratification for cancer patients based on transcriptomic data, using datasets from 25 cancer types in the TCGA database. To avoid cross-cancer data leakage and ensure test-set independence, each cancer-type dataset was divided into a training set (70%) and a validation set (30%) using stratified random sampling by cancer type. A multi-task joint-training strategy was employed to simultaneously fine-tune the RNABag model on datasets from 25 cancer types, thereby constructing a survival prediction model consisting of the RNABag encoder and a Cox proportional hazards decoding layer. The decoding layer used Cox partial-likelihood loss as the optimization objective, effectively handling the inherent time dimension and censored nature of survival data to achieve end-to-end modeling of patient survival risk. The C-index was used as the core evaluation metric to quantitatively measure the consistency between the risk scores predicted by the model and the actual survival ordering of patients. A C-index closer to 1 indicates stronger survival-risk stratification capability.

#### 1-Year Recurrence Risk Prediction

The 1-year recurrence risk prediction task was explicitly defined as a binary classification task to predict the risk of disease recurrence within 12 months after initial treatment in pan-cancer patients, aiming to identify populations at high risk of short-term recurrence and provide data support for determining follow-up intensity and early intervention strategies. This task used datasets from 31 cancer types in the TCGA database, with each cancer-type dataset divided into training and test sets using stratified random sampling by cancer type to ensure a reasonable data distribution and reproducibility of evaluation results. The RNABag model was fine-tuned within a multi-task learning framework, with the pre-trained encoder connected to a binary classification MLP decoder that learned discriminative features related to short-term recurrence through nonlinear mapping and output the probability of recurrence within 1 year via a sigmoid activation function. The model was optimized using cross-entropy loss to minimize the difference between predicted recurrence probability and true recurrence labels (0/1). AUROC was selected as the core evaluation metric to quantify the model’s overall ability to identify populations at high risk of short-term recurrence across 31 cancer types.

#### Liquid Biopsy Analysis

Liquid biopsy analysis aimed to validate the potential application of the RNABag model in non-invasive cancer detection and comprised three subtasks: plasma cfRNA cancer detection, TEP RNA-seq-based cancer detection, and TEP RNA-seq-based tumor localization. All liquid biopsy datasets were first subjected to standardized batch-effect correction to eliminate systematic variation arising from different sequencing platforms and research centers before being used for model training and validation, thereby ensuring the reliability of the experimental results.

#### Plasma cfRNA Cancer Detection

Transfer learning was conducted for plasma cfRNA cancer detection based on the tissue-sample cancer detection model. Specifically, the embedding module and encoder from the tissue-sample cancer detection model were retained and fine-tuned to adapt to the inherent characteristics of plasma cfRNA—fragmentation, low abundance, and high noise. An MLP classification decoder optimized for plasma signals was attached to the end of the model, and the unfrozen model parameters were updated using cross-entropy loss. Given the limited sample size of the plasma cfRNA datasets, five-fold cross-validation was employed for training to achieve robust fine-tuning and reduce the risk of overfitting. AUROC and precision were used as the core evaluation metrics, with precision specifically assessing the model’s ability to reduce false positives in clinical screening scenarios, in line with the practical requirements of non-invasive screening.

#### TEP RNA-seq-Based Cancer Detection

Transfer learning was conducted for TEP RNA-seq-based cancer detection based on the tissue-sample cancer detection model. Specifically, the embedding module and encoder from the tissue-sample cancer detection model were retained and fine-tuned to adapt to the characteristic expression patterns of TEP RNA-seq data. An MLP classification decoder was attached to the end of the model, and the unfrozen model parameters were updated using cross-entropy loss. To rigorously evaluate cross-dataset generalization, a single-dataset fine-tuning plus dual-dataset blind-testing strategy was employed. One of the three independent TEP RNA-seq datasets was selected as the fine-tuning set, whereas the other two datasets were kept completely isolated throughout training and model selection and used as independent test sets for blind testing of the fine-tuned model. This setup simulated the prediction stability of the model when faced with data from unknown research centers, different geographic origins, and various sequencing platforms in real-world clinical applications.

AUROC and precision were calculated on two independent external test sets to analyze the model’s generalization bias. The RNABag model was compared with mainstream models to quantitatively evaluate its robustness and advantages in scenarios involving non-tissue-derived TEP RNA-seq data.

#### TEP RNA-seq-Based Tumor Localization

The TEP RNA-seq-based tumor localization task aimed to validate whether the model could achieve pan-cancer tumor primary-site tracing based on TEP transcriptomic signals, thereby providing a potential non-invasive tool for the diagnosis and management of cancers of unknown primary. This task continued the strict single-dataset fine-tuning plus dual-dataset blind-testing strategy and established stringent inclusion criteria: only cancer types with no fewer than 100 samples in the fine-tuning set were retained to reduce model bias caused by extreme class imbalance.

The TEP RNA-seq-based tumor localization model was built on the embedding module and encoder of the pre-trained RNABag model, and both components were kept unfrozen during fine-tuning. This design enabled the model to adapt to the unique signal-distribution characteristics of TEP RNA-seq data and extract key information about tumor origin. The 128-dimensional feature vector output by the encoder was fed into a multiclass MLP decoder, with the output dimension matching the number of included cancer types. The model was optimized by minimizing cross-entropy loss to achieve multiclass classification of tumor primary sites, thereby mapping complex transcriptomic variations induced in TEPs back to specific tumor origins.

Confusion matrices were used to evaluate the localization accuracy of the model, intuitively displaying recognition accuracy for the tumor primary site of each cancer type and the misclassification patterns caused by transcriptomic similarities among different anatomical sites. By comparing the consistency of model performance in confusion matrices from the training set and independent test sets, the robustness of the RNABag model when faced with heterogeneous real-world TEP RNA-seq data was quantified, further validating the technical feasibility of pan-cancer localization based on TEP signals.

### Model Interpretation

#### SHAP-based Gene Attribution Analysis

To elucidate the biological mechanisms underlying the prediction decisions of the RNABag model and clarify the contribution of individual genes to model outputs, gradient-based SHAP were employed for gene attribution analysis, thereby transforming the model from a “black box” into an interpretable framework. The background distribution for SHAP analysis was constructed in the original gene-expression representation space to ensure consistency between the interpretation framework and the model input features. Given the large scale of the TCGA-GTEx merged dataset, k-means clustering (k = 500) was performed on all samples, with cluster centers used as background samples for SHAP estimation, effectively approximating the global data distribution while substantially reducing the computational cost of attribution analysis. For each tissue type or cancer type involved in downstream tasks, samples were randomly selected from the non-background data according to class-balance principles to ensure the statistical significance of SHAP value calculations and avoid analytical bias due to insufficient sample size or uneven distribution.

The SHAP value for each gene was calculated to quantify its contribution to the model’s predictions. First, the baseline output of the model was estimated using the cluster centers of the background samples. Then, the difference between the model’s predicted output for target samples and the baseline output was quantitatively attributed to each gene in the sample to obtain the SHAP value for each individual gene. For all samples of the same tissue type or cancer type, the mean absolute SHAP value for each gene was calculated to measure that gene’s importance to model decisions. Genes were ranked in descending order of mean SHAP value to identify core genes that played key roles in the model’s predictions.

#### Functional Enrichment Analysis

To investigate the biological significance of the core genes identified through SHAP-based gene attribution analysis and reveal the biological pathways and molecular mechanisms underlying the model’s predictions, functional enrichment analysis was performed on the 100 core genes with the highest mean SHAP values for each tissue type or cancer type. Using the GTEx database gene set as the reference database, two classical enrichment analysis methods were employed: Gene Ontology (GO)^49^ functional annotation and Kyoto Encyclopedia of Genes and Genomes (KEGG)^50^ pathway enrichment analysis. GO functional annotation characterized core genes across three dimensions—biological process, cellular component, and molecular function—thereby clarifying their basic biological roles. KEGG pathway enrichment analysis explored the key signaling pathways involving these core genes, revealing molecular regulatory networks related to tissue characteristics, cancer development, survival prognosis, and tumor origin. To control false positives caused by multiple hypothesis testing, the Benjamini-Hochberg^51^ method was used to adjust the P values of the enrichment results, and pathways with adjusted P values < 0.05 were defined as significantly enriched, thereby ensuring the reliability of the enrichment analysis.

## Data availability

TCGA and GTEx transcriptomic data used for model pretraining were obtained via recount3 (https://rna.recount.bio/). All independent validation datasets are publicly available. Solid tissue bulk RNA-seq datasets were accessed from the Gene Expression Omnibus (GEO) under accession numbers GSE140343^52^, GSE226069^53^, GSE179730^54^, GSE234304^55^, GSE103001^56^, GSE120795^57,58^, GSE126848^59^, GSE184336^60^, GSE85465^61^, GSE152415^62^, GSE119794^63^, GSE171485^64^, GSE130688^65^, GSE164665^66^, GSE196009^67^, GSE211398^68^, GSE133624^69^, GSE244957^70^, GSE66539^71^, GSE84776^72^, GSE229509^73^ and GSE151666^74^, and from cBioPortal (https://www.cbioportal.org/). Plasma cfRNA datasets are available under GEO accession numbers GSE174302^37^, GSE182824^38^, GSE216561^39^, GSE186587^40^ and GSE142987^41^. Tumor-educated platelet datasets are available under GEO accession numbers GSE68086^14^, GSE183635^15^ and GSE207586^42^. In-house clinical samples from the Affiliated Hospital of Wenzhou Medical University and the Affiliated Hospital of Xiangya School of Medicine are subject to institutional review board restrictions and are available from the corresponding author upon reasonable request with appropriate institutional approval. All other data are available from the corresponding author upon reasonable request.

---

**Supplementary Figure. 1.**
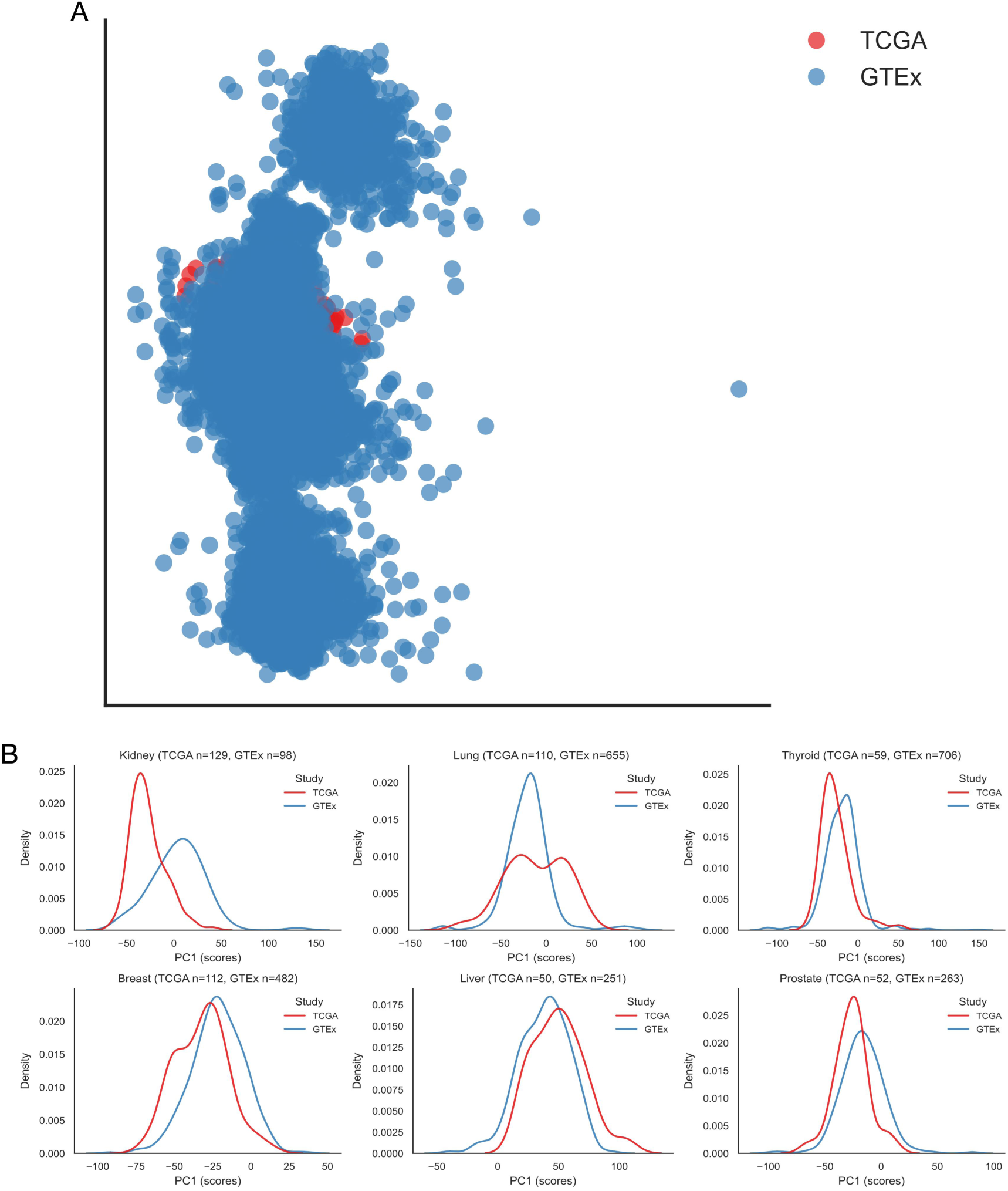
| Integration consistency and distribution characteristics of the TCGA and GTEx datasets. A, Two-dimensional projection of the global transcriptomic landscape. The UMAP scatter plot displays the overall expression distribution of samples from the TCGA (red) and GTEx (blue) databases. The two datasets exhibit a high degree of overlap and continuity in the reduced-dimensional space, indicating that systematic batch effects have been effectively eliminated or mitigated prior to downstream analysis. B, Expression distribution properties of specific organs. Kernel density estimation (KDE) plots depict the score distributions of kidney, lung, thyroid, breast, liver, and prostate samples on the first principal component (PC1). The distribution trajectories of the TCGA tumor cohorts (red) and the GTEx normal control cohorts (blue) along the principal component axis demonstrate biological consistency, providing a robust data foundation for cross-cohort cancer detection and tissue-of-origin tracing analysis.

**Supplementary Figure. 2.**
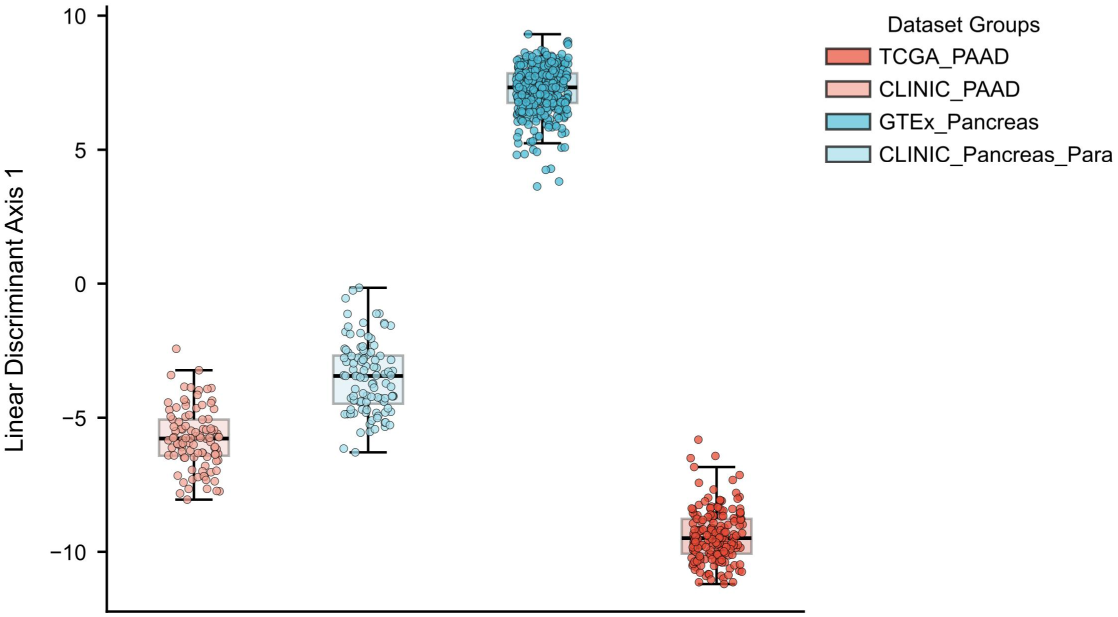
| Distribution of LDA1 scores across TCGA_PAAD, CLINIC_PAAD, GTEx_Pancreas, and CLINIC_Pancreas_Para samples. Boxplots represent the median and interquartile range, while overlaid points indicate individual samples. Significant differences between selected group pairs were assessed using two-sided Mann – Whitney U tests. ***P < 0.001.

